# Quantification of local matrix deposition during muscle stem cell activation using engineered hydrogels

**DOI:** 10.1101/2024.01.20.576326

**Authors:** Pamela Duran, Benjamin A. Yang, Eleanor Plaster, Madeline Eiken, Claudia Loebel, Carlos A. Aguilar

## Abstract

Adult stem cells occupy a niche that contributes to their function, but how stem cells remodel their microenvironment remains an open-ended question. Herein, biomaterials-based systems and metabolic labeling were utilized to evaluate how skeletal muscle stem cells deposit extracellular matrix. Muscle stem cells and committed myoblasts were observed to generate less nascent matrix than muscle resident fibro-adipogenic progenitors. When cultured on substrates that matched the stiffness of physiological uninjured and injured muscles, the increased nascent matrix deposition was associated with stem cell activation. Reducing the ability to deposit nascent matrix in muscle stem cells attenuated function and mimicked impairments observed from muscle stem cells isolated from old aged muscles, which could be rescued with therapeutic supplementation of insulin-like growth factors. These results highlight how nascent matrix production is critical for maintaining healthy stem cell function.

## Introduction

Age-related diseases manifest in nearly every tissue in the body such as skeletal muscle (sarcopenia), joint tissue (osteoarthritis), brain (Alzheimer’s) and a critical contributor of these pathologies is the loss of adult stem cell function.^[1]^ The functionality of adult stem cells is balanced between intrinsic (chromatin packaging and gene expression) and extrinsic mechanisms (interactions with the extracellular matrix or ECM).^[2, 3]^ An excellent example of age-associated stem cell dysfunction occurs in skeletal muscle, where muscle stem cells (MuSCs) undergo increases in chromatin accessibility^[4]^ and genomic interactions ^[5, 6]^ that drive changes in gene expression.^[7, 8]^ The intrinsic changes in MuSCs during aging are also associated with alterations^[9, 10]^ in the ECM.^[11, 12]^ Old aged skeletal muscle has been shown to lose fibronectin^[13]^, as well as displays increased stiffness^[14]^ or loss of elasticity^[15]^ from increases in total collagen content^[16]^ and changes in collagen type (increases in Collagen I, III and IV and reductions in collagen VI^[17]^). The change in the composition and mechanical properties of the ECM also coincides with reduced adhesion of MuSCs,^[18]^ that in turn promotes constitutive activation and eventual exhaustion.^[19, 20]^ Yet, linking changes in the ECM^[21]^ that occur in aging has been hampered by a lack of knowledge of how MuSCs directly contribute to their niche.

Biomaterials are excellent tools to investigate the influence of biochemical and biophysical cues of MuSCs. Synthetic materials composed of polyethylene glycol (PEG) functionalized with an adhesive peptide or ECM component have been used to mimic native muscle^[22–25]^ and showed that MuSCs cultured on a two-dimensional substrate similar to the stiffness of a healthy muscle^[23]^ promoted a quiescent phenotype. PEG-based materials functionalized with peptides that replicate fibronectin and change stiffness in response to light also showed MuSC culture on 2D substrates that soften in situ from 11kPa to 6 kPa, as observed after injury, led to increased proliferation and differentiation.^[22]^ PEG-based materials functionalized with laminin or fibronectin mimetics also showed that mechanically stiffer hydrogels (elastic moduli ∼40 kPa) were associated with lower proliferation and dysfunctional activation.^[24]^ Biomaterials thus offer the unique ability to tune material properties to replicate ECM^[26]^ and aspects of the MuSC microenvironment.^[27–30]^ However, how MuSCs produce and remodel their niche after injury and aging has not been evaluated.

Herein, we utilized biomaterials-based systems and metabolic labeling^[31]^ to evaluate how MuSCs deposit and remodel their matrix. We first compared nascent matrix deposition between MuSCs, myoblasts and fibro-adipogenic progenitors (FAPs). We observed that both types of myogenic cells produce less matrix than FAPs. We then cultured MuSCs on substrates that mimic physiological muscle stiffness during homeostasis and observe increased nascent matrix deposition as MuSCs activate and exit quiescence. When MuSCs were cultured on softer matrices as observed during early muscle regeneration, we found higher MuSC activation and increases in matrix deposition. We next found that blocking the ability to deposit new matrix attenuated MuSC fusion, in line with previous observations that niche remodeling is a critical determinant of successful regeneration.^[32]^ Given aging has been demonstrated to induce impairments in MuSCs, we quantified how MuSCs isolated from different stages of life deposit new matrix. We detected significant reductions in nascent matrix deposition for old aged MuSCs when compared to young, which was rescued with insulin-like growth factor stimulation. Overall, this study expands our understanding of MuSCs and how adult stem cells remodel their niche.

## Results and Discussion

### Metabolic labeling of secreted matrix from muscle stem cells and fibro-adipogenic progenitors

Skeletal muscle contains a population of MuSCs that are responsible for muscle repair after injury and mesenchymal progenitors (fibro-adipogenic progenitors or FAPs) that signal to MuSCs in their niche.^[33]^ To begin to understand how MuSCs and FAPs deposit new matrix, we first isolated MuSCs^[34]^ and FAPs from uninjured hindlimb muscles of two types of young transgenic mice (3 months). The first model harbors a red fluorescent reporter (TdTomato) in Pax7 (*Pax7^CreERT2^-Rosa26^TdTomato^*), and the second mouse model contains a TdTomato reporter in the PDGFRα locus (PDGFRα^Cre^-Rosa26^Tdtomato^), respectively. Both freshly isolated MuSCs and FAPs were cultured on matrigel in media containing l-azidohomoalanine (AHA), which is a reactive noncanonical methionine analog that is incorporated into newly synthesized proteins. An azido moiety in AHA facilitates detection with a biotinylated alkyne and imaging of newly synthesized extracellular proteins (**Fig. 1A**). We fixed and imaged MuSCs and FAPs after 5 days of culture, and observed both cell types deposit nascent matrix as puncta (**Fig. 1B**). FAPs were observed to generate more matrix than MuSCs (n=3, p=0.03, one-way ANOVA with Tukey’s range test), which is in line with previous observations.^[11]^ We next asked if the observed puncta overlapped with focal adhesions to glean if cells were depositing matrix to strengthen contacts with the underlying substrate, and stained cells for vinculin (**Supp. Fig. 1A-B**). However, we detected minimum overlap between vinculin and nascent matrix suggesting a weak attachment of the cells on a two-dimensional culture system at the time point analyzed. To determine if the nature of nascent protein deposition into punta was unique to primary cells isolated from skeletal muscle, we contrasted C2C12 myoblasts in the same media containing AHA. We detected comparable levels of nascent protein deposition between MuSCs and C2C12s, both of which were lower than FAPs (n=3, p=0.01, one-way ANOVA with Tukey’s range test, **Figs. 1B-C**). These results demonstrate we can track nascent matrix deposition from skeletal muscle cells in an *in vitro* culture system.

**Figure 1.**
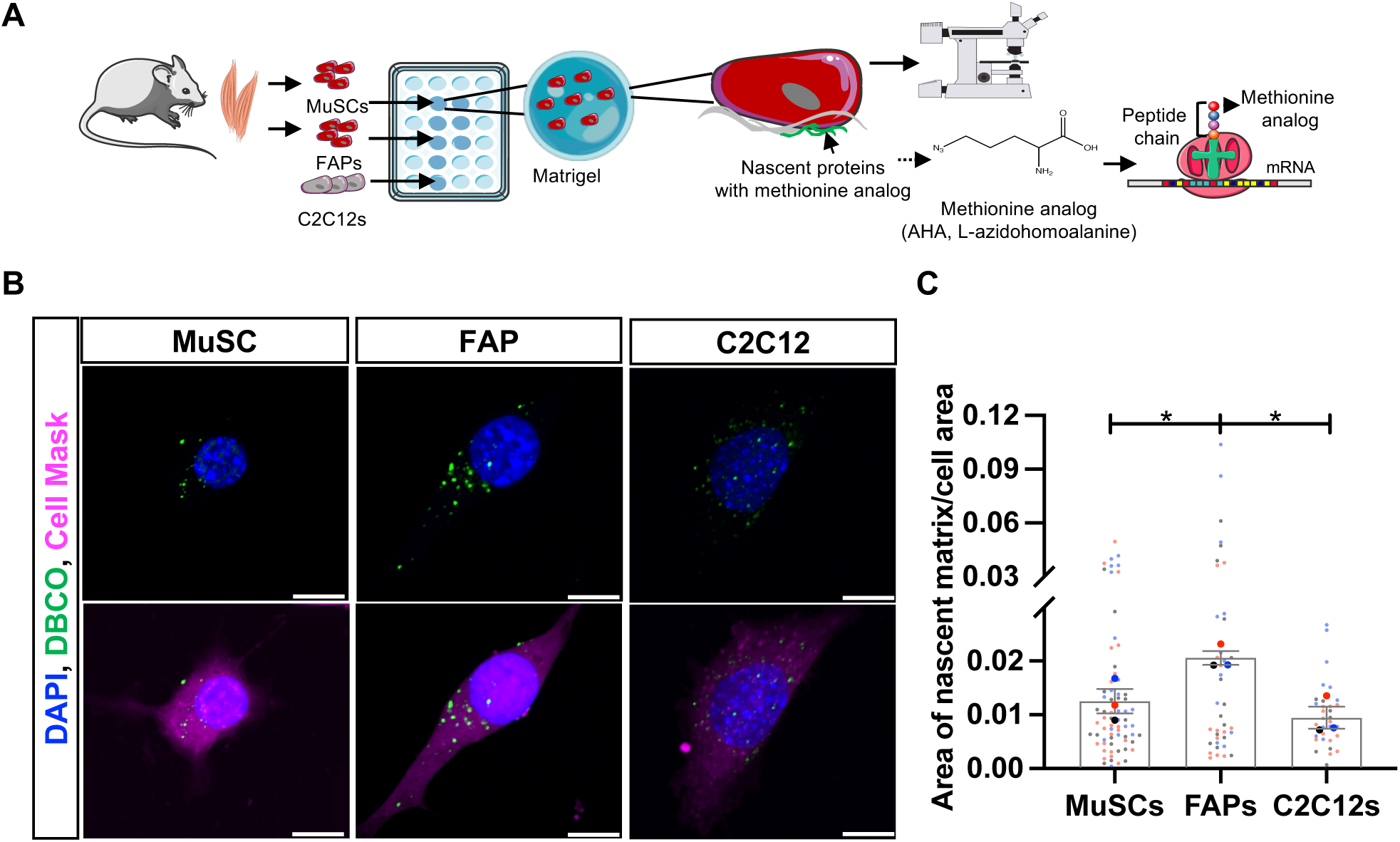
Muscle stem cells (MuSCs) and myoblasts deposit less nascent matrix than fibro-adipogenic progenitors (FAPs). **A)** Muscle stem cells (MuSCs), fibro-adipogenic progenitors (FAPs), and C2C12 myoblasts were cultured in methionine analog-based media to visualize nascent proteins. **B)** Representative images of the different cell types cultured methionine analog-based media for 5 days. Blue: DAPI, Green: Nascent Matrix, Pink: Cell Mask. Scale = 10 !m. **C)** Quantification of the area of nascent matrix normalized to cell area. p-values derived from one-way ANOVA followed by pairwise comparisons with Tukey’s range test. *p<0.05; mean ± SEM; small circles represent single cells (n=10-20/replicate) and large circles indicate biological replicates or experimental runs (n=3 animals).

### Nascent matrix deposition is associated with activated muscle stem cells

MuSCs have been shown to be highly sensitive to physiological conditions, and exhibit enhancements in quiescence when cultured ex vivo on substrates that mimic the stiffness of uninjured skeletal muscle. To glean whether culture of MuSCs on physiologically matched mechanical properties of skeletal muscle (elastic modulus of ∼11 kPa) altered the ability to deposit nascent matrix, we created covalently crosslinked hydrogels composed of hyaluronic acid (HA) functionalized with norbornene groups (Nor-HA, **Fig. 2A**) and modified the HA with the cell-adhesive peptide arginylglycylaspartic acid (RGD). We confirmed the elastic modulus of the NorHA hydrogels by rheology (**Supp. Fig. 2A**) and tracked nascent matrix deposition from MuSCs over a time course (1, 3 and 5 days, **Fig. 2B**). We observed increases in nascent matrix deposition at five days in culture when compared to one or three days (n=2-4 biological replicates, 5 vs 1 day p=0.004, 5 vs 3 days p=0.005, one-way ANOVA with Tukey’s range test, **Fig. 2C**), respectively. To determine state of the MuSCs, we immunostained for Pax7 and MyoD and co-stained for nascent matrix as well as DAPI (**Fig. 2D**). We found that MuSCs that increased MyoD also displayed increased nascent matrix deposition, when compared to MuSCs that displayed lower expression of MyoD (n=3 biological replicates, p=0.02, two-sided student’s t-test, **Fig. 2E, Supp. Fig. 2B**). These results suggest that activated MuSCs secrete more matrix. To further determine if quiescent MuSCs produced less nascent matrix than activated MuSCs, we immunostained for DEAD-Box Helicase 6 (DDX6), an RNA helicase that is enriched in stress granules and is strongly associated with MuSC quiescence, and nascent matrix (**Figs. 2F-G**). We observed that MuSCs that contained higher expression of DDX6 also produced less nascent matrix, when compared to MuSCs that displayed less DDX6 and generated more nascent matrix (n=3 biological replicates, p=0.03, two-sided student’s t-test, **Fig. 2G**).

**Figure 2.**
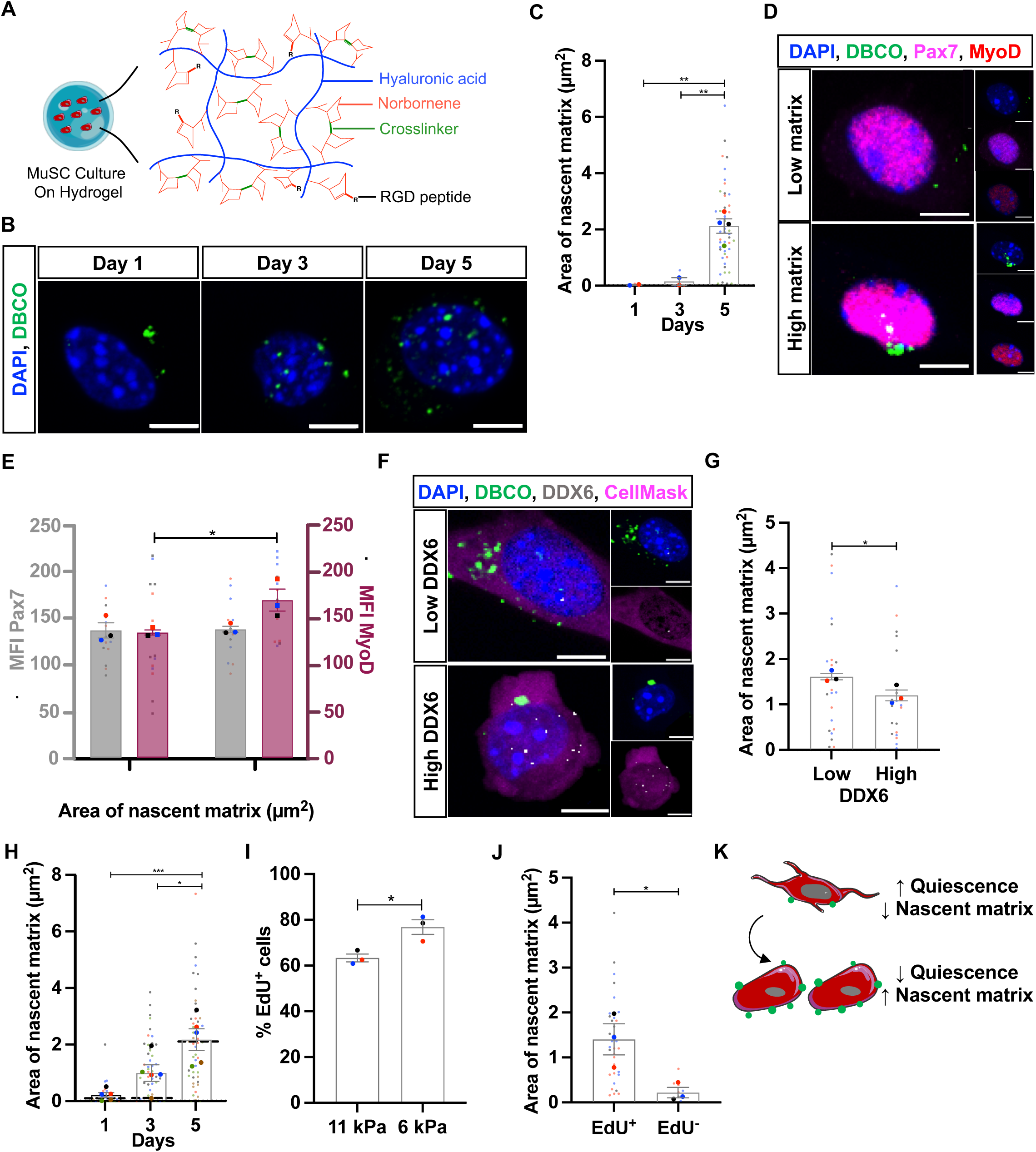
Increased nascent matrix deposition is associated with muscle stem cell activation. **A)** Muscle stem cells (MuSCs) were isolated and cultured on norbornene-modified hyaluronic acid (Nor-HA) hydrogels with physiological stiffness that replicate an uninjured muscle (11 kPa) or injured muscle during regeneration (6 kPa). **B)** Representative images of MuSCs cultured on 11kPa Nor-HA hydrogels for up to 5 days. Blue: DAPI, Green: Nascent Matrix. Scale = 5 !m. **C)** Quantification of nascent matrix deposition from images in B). Single cells=10-25/replicate, biological replicates=2-4. **D)** Representative images of MuSCs cultured on Nor-HA hydrogels for 5 days show two categories: cells that produce low amounts of nascent matrix and cells that produce high amounts of nascent matrix. Blue: DAPI, Green: Nascent Matrix, Pink: Pax7, Red: MyoD. Scale = 5 !m. **E)** Quantification of D) on single cells that were partitioned by low and high amounts of nascent matrix deposition plotted against the mean fluorescence intensity (MFI) of Pax7 and MyoD. Single cells=10/replicate, biological replicates=3. **F)** Representative images of MuSCs cultured for 5 days demonstrating nascent proteins, and DDX6 expression. Blue: DAPI, Green: Nascent Matrix, Pink: Cell Mask, Gray: DDX6. **G)** The number of DDX6+

Since injured skeletal muscle has been shown to decrease elasticity several days after injury when MuSCs are activating and beginning to differentiate, ^[22]^ we created Nor-HA hydrogels with 6 kPa stiffness by adjusting the concentration of the cross-linker (**Supp. Fig. 2A**). We cultured freshly isolated MuSCs over a time course as above and observed similar amounts of nascent matrix deposition for MuSCs cultured on 6 kPa when compared to 11 kPa hydrogels at 1 and 5 days (n=5 biological replicates, 5 vs 1 day p=0.001, 5 vs 3 days p=0.03, one-way ANOVA with Tukey’s range test, **Fig. 2H**). However, at 3 days, a relative increase in nascent matrix deposition was observed for MuSCs cultured on the 6 kPa hydrogel when compared to 11 kPa hydrogels (**Fig. 2H**), respectively. We reasoned that the increase in the kinetics of MuSC nascent matrix deposition was due to increased activation of MuSCs on the softer substrates. To determine if MuSCs cultured on 6 kPa substrates displayed increased activation compared to MuSCs cultured on 11 kPa substrates, we added EdU to the culture medium and examined changes in DNA synthesis for the two conditions. In line with our previous observations, we detected a significant increase in EdU incorporation for MuSCs cultured on 6 kPa substrates when compared to 11 kPa (n=3 biological replicates, p=0.02, two-sided student’s t-test, **Fig. 2I**). Furthermore, EdU^+^ MuSCs displayed an 84.6% increase in nascent matrix deposition when compared to EdU^-^ MuSCs (n=3 biological replicates, p=0.03, two-sided student’s t-test, **Fig. 2J**). These results demonstrate that MuSCs increase nascent matrix deposition as they activate and break quiescence.

To further elucidate the matrix composition of MuSCs during activation, we analyzed previously obtained RNA-sequencing data^[4]^ of freshly isolated MuSCs in uninjured and injured muscle after barium chloride injection. Following differential expression analysis, we segregated the genes related to the core matrisome based on a previous classification^[31, 35]^ and the data were visualized with a heatmap (**Supp. Fig. 2C**). Few genes (7 genes) were significantly differentially expressed (DE) at 1-day post injury (dpi) with the majority being changed at 3 dpi (with 117 genes) and decreased at 7 dpi (29 DE genes). ECM remodeling started at 3 dpi with downregulation of 85% of the DE genes, which corresponded mostly to glycoproteins. From the upregulated genes at each of the timepoints, we then obtained the proportion of the genes that corresponded to the core matrisome categories (**Supp. Fig. 2D**). At 1 dpi, the genes were related to collagens [*Collagen type VXIII alpha 1* (Col18a1), *Collagen type XXVII alpha 1* (Col27a1), *Collagen type XXVIII alpha 1* (Col28a1)] and glycoproteins [*Dentin Matrix Acidic Phosphoprotein 1* (Dmp1), *Leucine-rich repeat LGI family member 4* (Lgi4), *Tenascin N* (Tnn)]. The upregulation of genes corresponding to glycoproteins [*Otogelin* (Otog), *Insulin-like growth factor binding protein 2* (Igfbp2), Dmp1, Slit1, *Fibronectin type III domain containing 7* (Fndc7), *Nephronectin* (Npnt), *Acetylcholinesterase-Associated Collagen* (Colq), *Cellular communication network 4* (Wisp1), *Insulin-like growth factor binding protein 5* (Igfbp5), *laminin b1* (Lamb1), *Von Willebrand Factor A Domain Containing 9* (Vwa9)] predominated at 3 dpi with upregulation of few collagens [*Collagen type XIII alpha 1* (Col13a1), Col18a1, *Collagen type XXVI alpha 1* (Col26a1)] and proteoglycans [*Podocan like 1* (Podnl1)] genes. The pattern continued at the later time point with the majority of the genes related to glycoproteins [Otog, Slit1, Dmp1, *Von Willebrand Factor C* (Vwce), *Matrillin-4* (Matn4), *Periostin* (Postn), *Nidogen 2* (Nid2), Lgi4, Wisp1, Lamb1] and some collagens [*Collagen type III alpha 1* (Col3a1), *Collagen type VI alpha 2* (Col6a2), Col18a1, *Collagen type XXVI alpha 1* (Col26a1), Col27a1]. Overall, these results indicate that MuSCs upregulate genes during activation mostly related to glycoproteins.^[3]^

### Perturbation of nascent matrix deposition alters the functionality of muscle stem cells

To evaluate if changes in the functionality of MuSCs are enacted with alterations to deposit nascent matrix, we compared MuSCs treated with a small molecule that interrupts the trafficking and secretion of extracellular proteins between the endoplasmic reticulum and golgi apparatus^[36, 37]^ (2- (4-fluorobenzoylamino)-benzoic acid methyl ester or Exo-1) and untreated controls (**Fig. 3A**). Exo-1 treated MuSCs exhibited significant reductions in nascent matrix deposition (n=3 biological replicates, p=0.04, two-sided student’s t-test, **Fig. 3B**), and comparable levels of proliferation (n=4 biological replicates, p=0.1, two-sided student’s t-test, **Supp. Fig. 3A**). Differentiation and fusion of treated and control MuSCs into myoblasts and myotubes revealed reductions in fusion index (n=3 biological replicates, p=0.02, two-sided student’s t-test) and myotube width (n=3 biological replicates, p=0.01, two-sided student’s t-test) for Exo-1 treated MuSCs when compared to untreated controls, respectively (**Figs. 3C-E**). These results suggest nascent protein deposition and remodeling of the underlying niche is critical for MuSC activation and fusion into multinucleated myotubes.

**Figure 3.**
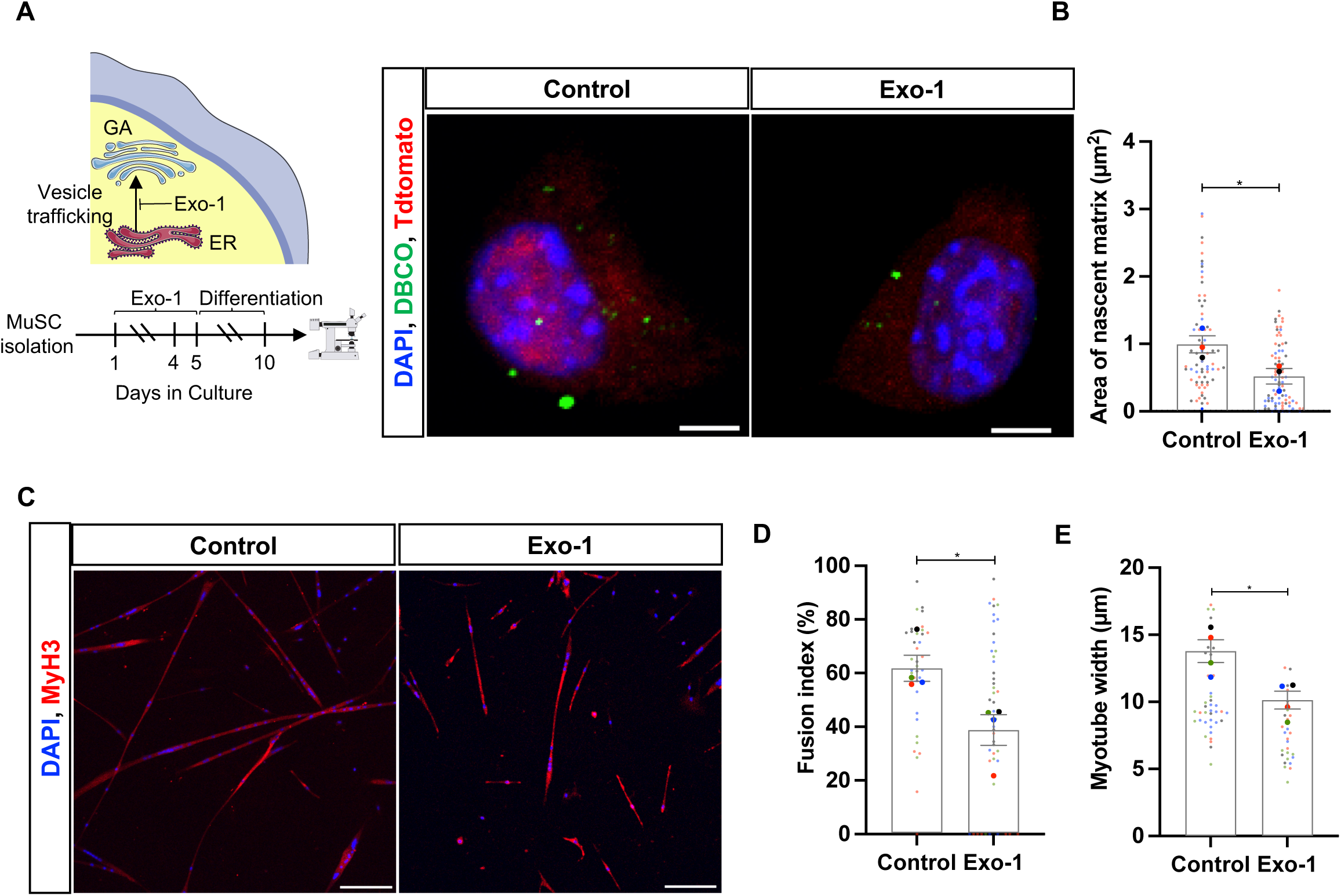
Nascent matrix deposition alters the functionality of muscle stem cells (MuSCs). **A)** MuSCs were isolated and incubated with Exo-1 to perturb the secretion of proteins (GA= Golgi Apparatus, ER= Endoplasmic Reticulum). Representative images indicate the nascent proteins after 5 days of culture. Scale bar = 5 µm. **B)** Quantification of the area of nascent matrix with and without Exo-1 at 5 days. Single cells= 20-30/replicate, biological replicates= 3. **C)** At 5 days of culture, MuSCs were incubated with differentiation media to induce fusion. Representative images of myosin heavy chain 3^+^ cells at 10 days of culture. Scale bar= 200 µm. Quantification of fusion index **(D)** and myotube width **(E)** with and without Exo-1. Myotubes= 20-30/replicate, biological replicates=4. p-values derived from student’s t-test. *p<0.05; mean ± SEM; small circles represent single cells and large circles indicate the biological replicates.

### Aging is associated with a progressive decrease in muscle stem cell (MuSC) nascent matrix deposition

MuSCs have been shown to exhibit dysfunction in old age^[38]^, and a loss of proteostasis and trafficking has been suggested to contribute to this pathological transformation.^[39]^ To investigate whether aging impinged on ability to deposit nascent matrix, MuSCs were isolated from hindlimb muscles across different ages (Young: 2-4 months, Middle-Aged: 13 months, Aged: 17-20 months, Old: 24-26 months, **Fig. 4A**) and cultured as above. The area of nascent matrix deposition was quantified over a time course (**Fig. 4B-C**). Both young and middle-aged MuSCs displayed an increase in nascent matrix at 5 days compared to 1 day, respectively (n=2-5 biological replicates, young-p<0.0001, middle-aged-p=0.029, two-way ANOVA with Tukey’s range test, **Fig. 4B**). However, no changes in matrix deposition between the time points were identified for aged and old MuSCs at day 5 and decreases in nascent matrix were observed for aged and old MuSCs when compared to young and middle-aged MuSCs, respectively (n=2-5 biological replicates, aged vs young-p=0.007, aged vs middle-aged-p=0.03, old vs young-p<0.0001, old vs middle-aged-p=0.002, two-way ANOVA with Tukey’s range test, **Fig. 4B-C**).

**Figure 4.**
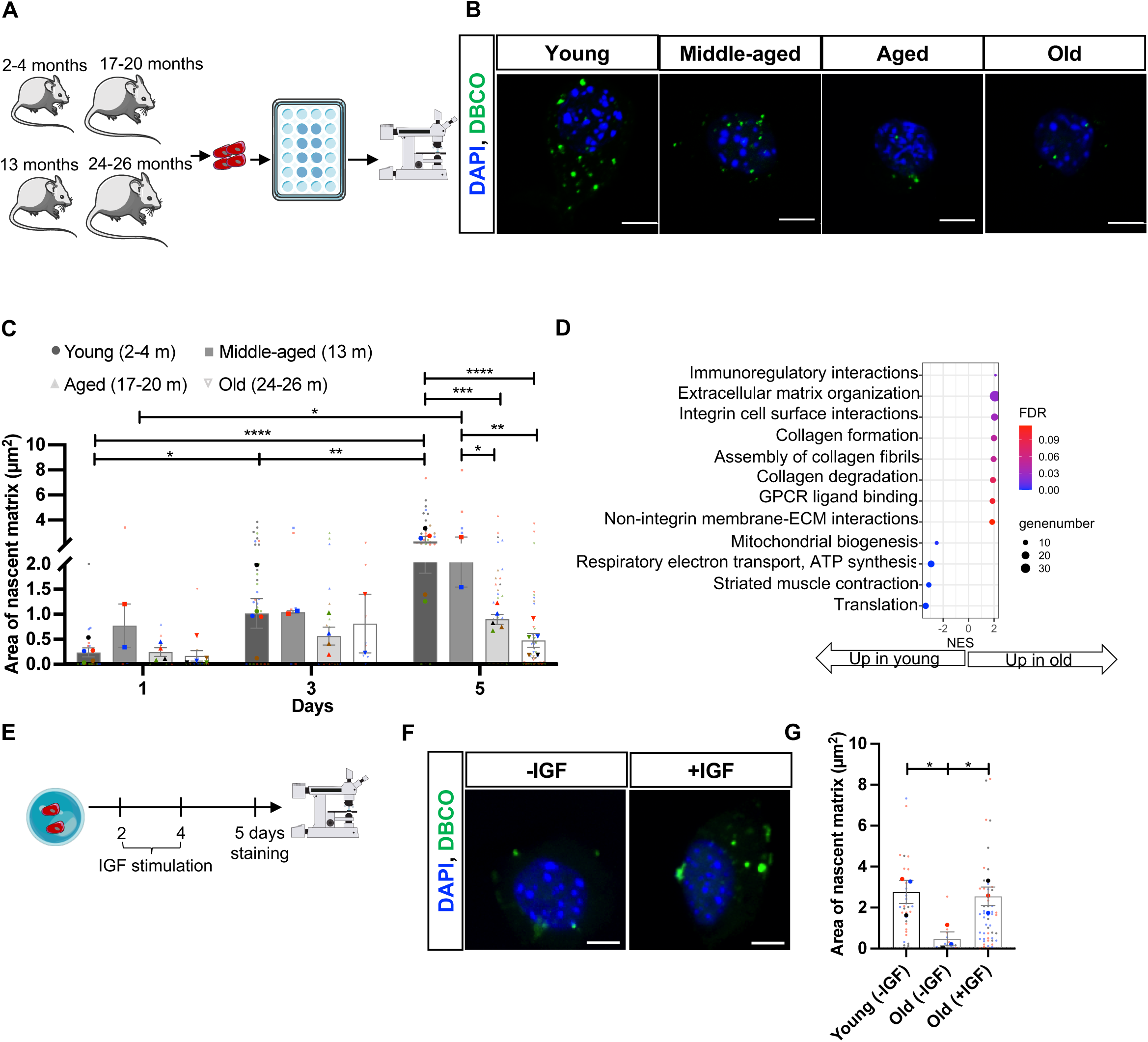
Aging is associated with a progressive decrease in muscle stem cell (MuSC) nascent matrix deposition. **A)** MuSCs were isolated from mice across different ages and cultured on the norbornene-modified hylauronic acid (Nor-HA) hydrogels for up to 5 days. **B)** Representative images of MuSCs (from young to old mice) at 5 days of culture. **C)** Quantification of the area of nascent matrix. Single cells= 10-25/replicate, biological replicates= 2-5. **D)** RNA was isolated from young and old MuSCs cultured for 5 days on Nor-HA hydrogels, prepared into sequencing libraries and sequenced. Gene set enrichment analysis of ranked significantly differentially expressed genes of old vs young groups (1532 genes) using the Reactome pathway database. Positive or negative normalized enrichment scores (NES) represent up- or downregulation, respectively. **E)** MuSCs from young and old mice were isolated, cultured on Nor-HA hydrogels and stimulated with IGF-1 (insulin like growth factor 1). **F)** Representative images of old MuSCs with and without IGF-1 treatment. **G)** Quantification of the area of nascent matrix across groups. Single cells= 10-25/replicate, biological replicates= 3. p-values derived from one- or two-way ANOVA followed by pairwise comparisons with Tukey’s range test. *p<0.05, **p<0.01, ***p<0.001, ****p<0.0001; mean ± SEM; small circles represent single cells and large circles indicate the biological replicates.

To gain insights into mechanisms potentially contributing to the impairment in nascent matrix, we performed RNA-sequencing on replicates of young and old MuSCs cultured for 5 days on the Nor-HA hydrogels. A strong agreement (Spearman coefficient=0.96) was observed for each biological replicate from MuSCs isolated from young and old groups (**Supp. Fig. 4A**). Differential expression analysis indicated that 1532 genes were changes in old vs young MuSCs. Pathway annotation using gene set enrichment analysis (GSEA)^[40]^ demonstrated increased expression of genes associated with pathways related to translation [*mitochondria ribosomal proteins* L42, S6, L34, L52, L55, *ribosomal proteins* S23, S13, S5, S15A, S18, S2, S3A1, *eukaryotic translation initiation factor 3, subunit I* (Eif3i)] and muscle contraction [*troponin I skeletal fast 2* (Tnni2), *troponin T1 skeletal slow* (Tnnt1), *troponin C slow skeletal* (Tnnc1), *actin alpha muscle 1* (Actc1), *myosin light polypeptide 4* (Myl4), *troponin C2 fast* (Tnnc2), *titin-cap* (Tcap), *myosin light polypeptide 1* (Myl1), *troponin I skeletal slow 1* (Tnni1), *nebulin* (Neb)] for young MuSCs, which was in contrast to old MuSCs that displayed increases in expression of genes related to collagen formation and degradation [*matrix metallopeptidase 9* (Mmp9), *matrix metallopeptidase 2* (Mmp2), *matrix metallopeptidase 12* (Mmp12), *collagen type IV alpha 1* (Col4a1), *collagen type IV alpha 2* (Col4a2), *collagen type IV alpha 5* (Col4a5), *collagen type V alpha 2* (Col5a2), *collagen type V alpha 3* (Col5a3), *collagen type VIII alpha 1* (Col8a1), *collagen type XIV alpha 1* (Col14a1), *collagen type XV alpha 1* (Col15a1), *collagen type XVI alpha 1* (Col16a1), *peroxidasin* (Pxdn), *lysyl oxidase* (Lox), *lysyl oxidase-like 4* (Loxl4)], and non-integrin and integrin interaction with the extracellular matrix [*integrin beta 2* (Itgb2), *intercellular adhesion molecule1* (Icam1), *collagen type IV alpha 5* (Col4a5), *collagen type XVI alpha 1* (Col16a1), *tenascin C* (Tnc), *collagen type V alpha 3* (Col5a3), *integrin alpha 2* (Itga2), *thrombospondin 1* (Thbs1), *integrin beta 3* (Itgb3), *collagen type IV alpha 1* (Col4a1), *collagen type VIII alpha 1* (Col8a1), *integrin alpha 5* (Itga5), *collagen type V alpha 2* (Col5a2), *F11 receptor* (F11r), *collagen type IV alpha 2* (Col4a2), *integrin beta 8* (Itgb8), *integrin alpha V* (Itgav), *syndecan 3* (Sdc3), *protein kinase C alpha* (Prkca)] (**Fig. 4D**). These results are consistent with previous observations that old MuSCs exhibit defects in protein synthesis and translational control.^[4, 41]^ Since targeting the insulin like growth factor 1 (IGF-1) pathway^[42]^ has been shown to stimulate protein phosphorylation and synthesis in skeletal muscle^[43]^ through phosphoinositide-3-kinase–protein kinase B/Akt (PI3K-PKB/Akt) signaling cascades,^[44]^ we treated old MuSCs with and without IGF-1. At 5 days of culture, the area of nascent matrix was quantified for young and old (with and without IGF-1) MuSCs (**Fig. 4E**). In line with our previous results, a decrease in nascent matrix was observed in old MuSCs compared to young MuSCs, respectively (n=3 biological replicates, p=0.03, one-way ANOVA with Tukey’s range test, **Fig. 4F-G**). However, for old MuSCs supplemented with IGF-1, we detected an increase in the area of nascent matrix and levels analogous with young MuSCs (p=0.94, **Fig. 4F-G**). These results demonstrate that old MuSCs display an impairment in nascent matrix deposition that can be rescued with IGF-1 stimulation.

## Conclusions

In this study, we utilized biomaterial-based systems to track nascent matrix deposition in stem cells and progenitors isolated from skeletal muscle. We first observed that FAPs display an increased area of nascent matrix when compared to MuSCs. We next detected that MuSCs deposit nascent matrix as they activate and exit quiescence, and this process was accelerated when MuSCs were cultured on substrates that mimicked physiological injured muscle stiffness. When MuSCs were cultured with a reversible inhibitor of exocytosis, Exo-1, and the secretion of extracellular matrix proteins was reduced, we detected a decrease in the fusion of MuSCs. Given old aged MuSCs have been shown to display defects in translation and niche remodeling, we demonstrated an impairment in nascent matrix deposition with aging, which could be restored with IGF-1 stimulation.

How stem cells and progenitors coordinate and maintain homeostasis in tissue remains an active area of research,^[45]^ and recent studies have suggested that cells divide labor^[46]^ and specialize to perform different functions. Tracking the deposition of nascent matrix proteins from MuSCs and FAPs showed FAPs deposit 66% more matrix than MuSCs. MuSCs and FAPs are proximal in tissue and critically rely on each other to enact appropriate regeneration. Our results suggest that FAPs may regulate matrix type and magnitude, and MuSCs respond to these cues through activation and differentiation. This relationship is in line with several studies^[9, 47]^ showing interruptions to FAP signaling and extracellular protein deposition result in MuSC dysfunction.^[48–50]^ While more studies are needed to further elucidate how FAPs shape and remodel the ECM and MuSCs correspondingly change state and respond, engineered platforms^[51]^ may offer a unique system^[52]^ to evaluate this relationship.

The microenvironment of MuSCs is a sensitive regulator of regenerative potential, but how MuSCs actively remodel their niche has been understudied. Our results show that as MuSCs activate and break quiescence, the cells deposit new matrix. We observed nascent matrix deposition as puncta, as opposed to filaments, which may be due to relatively weak attachment, which is in agreement with minimal overlap with immunostaining for focal adhesions. Furthermore, MuSCs during activation tend to be mobile^[53]^ and do not generate appreciable tractional forces in two-dimensional culture, which prevent extension and alignment of underlying nascent proteins. We speculate that myoblasts cultured in 3D systems^[54]^ may enhance matrix deposition and remodeling, as has been observed for differentiating neural stem cells,^[55]^ which also display minimal tensional forces in 2D.^[56]^ However, 3D hydrogels are usually synthesized with physical crosslinking to improve cell encapsulation and viability, which reduces the manipulation over the mechanical properties of the biomaterial.^[57]^ We observed that MuSC deposition of nascent proteins seemed to approach a threshold as they activate, as similar levels of nascent protein deposition was observed in myoblasts and when MuSCs were cultured on a less stiff substrate that mimicked muscle injury. Since perturbation of nascent protein deposition via Exo-1 treatment attenuated fusion but did not impinge on cellular growth / proliferation, we speculate extracellular nascent matrix deposition may be used to assess maintenance of quiescence^[58]^ and regenerative efficacy of MuSCs cultured in vitro.

MuSCs undergo intrinsic and extrinsic changes with aging, leading to alterations in their activation, self-renewal, and differentiation.^[38]^ We observed impairments in nascent matrix deposition for old MuSCs, similar to young MuSCs treated with Exo-1. We speculate that the reduced area of nascent matrix deposition may be due to the decoupling between the transcriptome and proteome in aged MuSCs,^[11]^ and the translational control.^[40]^ To further understand molecular drivers of defects in old aged MuSCs, we performed RNA-Seq and detected increased expression of genes related to collagen types, as well as downregulation of genes associated with translation, which is in line with previous observations.^[38, 39]^ Given IGF-1 has been shown to enhance protein synthesis and protect myoblasts from oxidative stress,^[59, 60]^ we found IGF-1 treatment restored nascent matrix deposition in old aged MuSCs back to levels of young MuSCs. We cannot exclude the possibility that the impairment in nascent matrix deposition observed in old aged MuSCs may also be derived from exacerbated ECM degradation. Our RNA-seq data showed upregulation of matrix metalloproteinases, which have previously been found to attenuate MuSC regenerative potential, ^[11, 61]^ and future studies will aim to further understand MuSC balance between nascent protein deposition and breakdown.

In this study, we generated a library of HA-based hydrogels and metabolic labeling strategies to investigate how MuSCs deposit nascent matrix. Future systems will aim to incorporate 3D encapsulation and co-culture with other skeletal muscle cells to provide insights into paracrine interactions that alter nascent matrix deposition of MuSCs.

## Materials and Methods

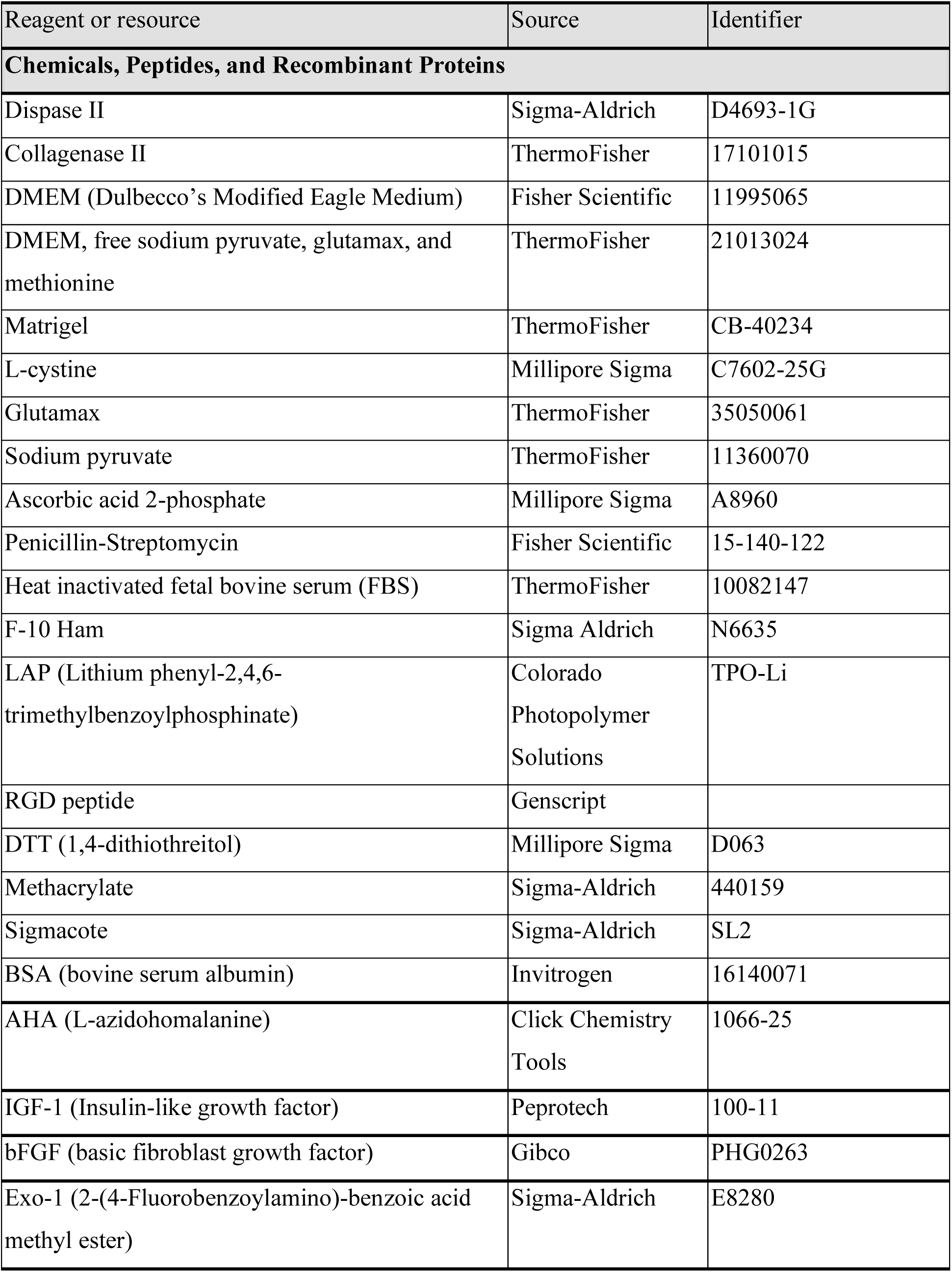
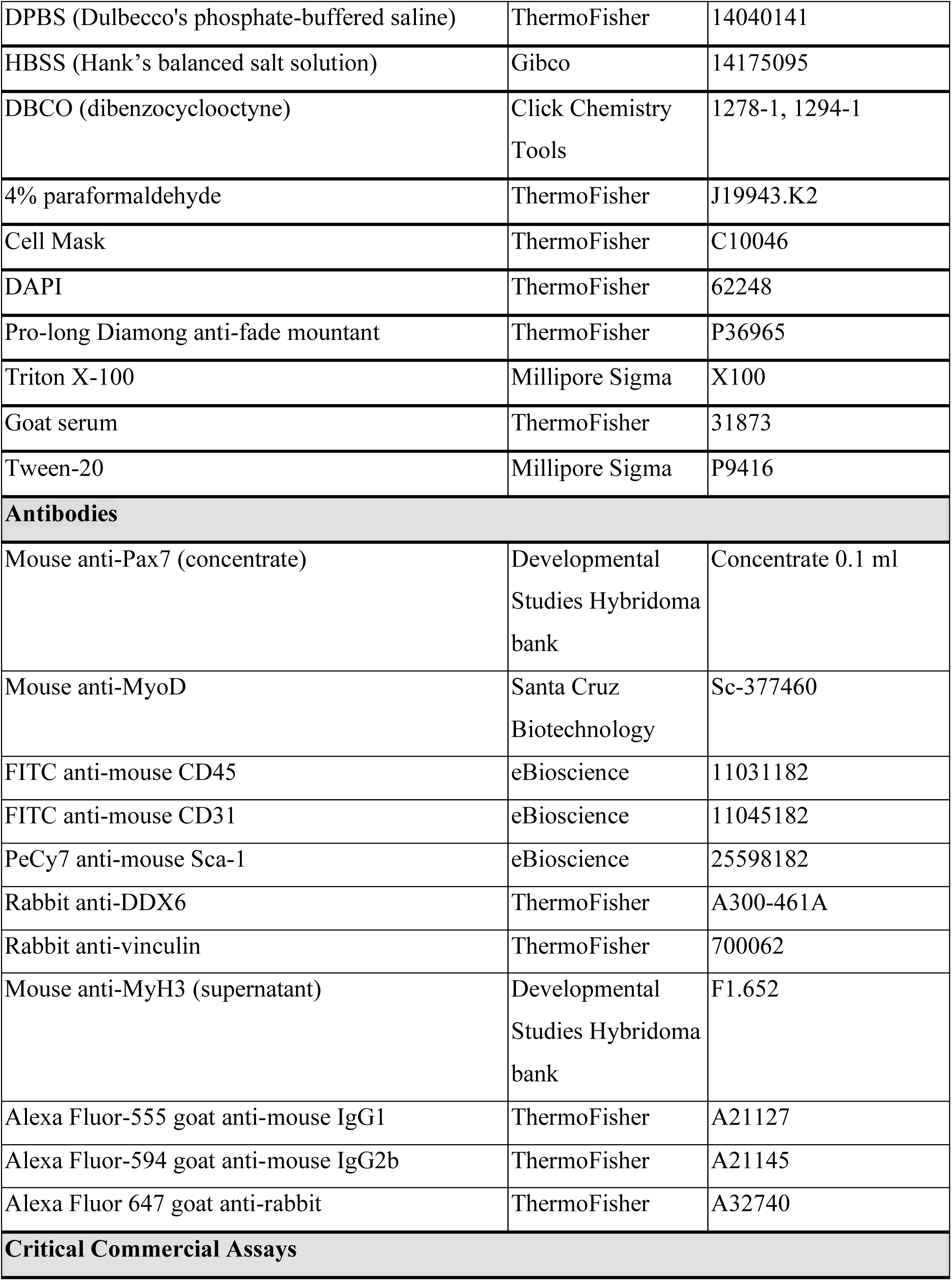
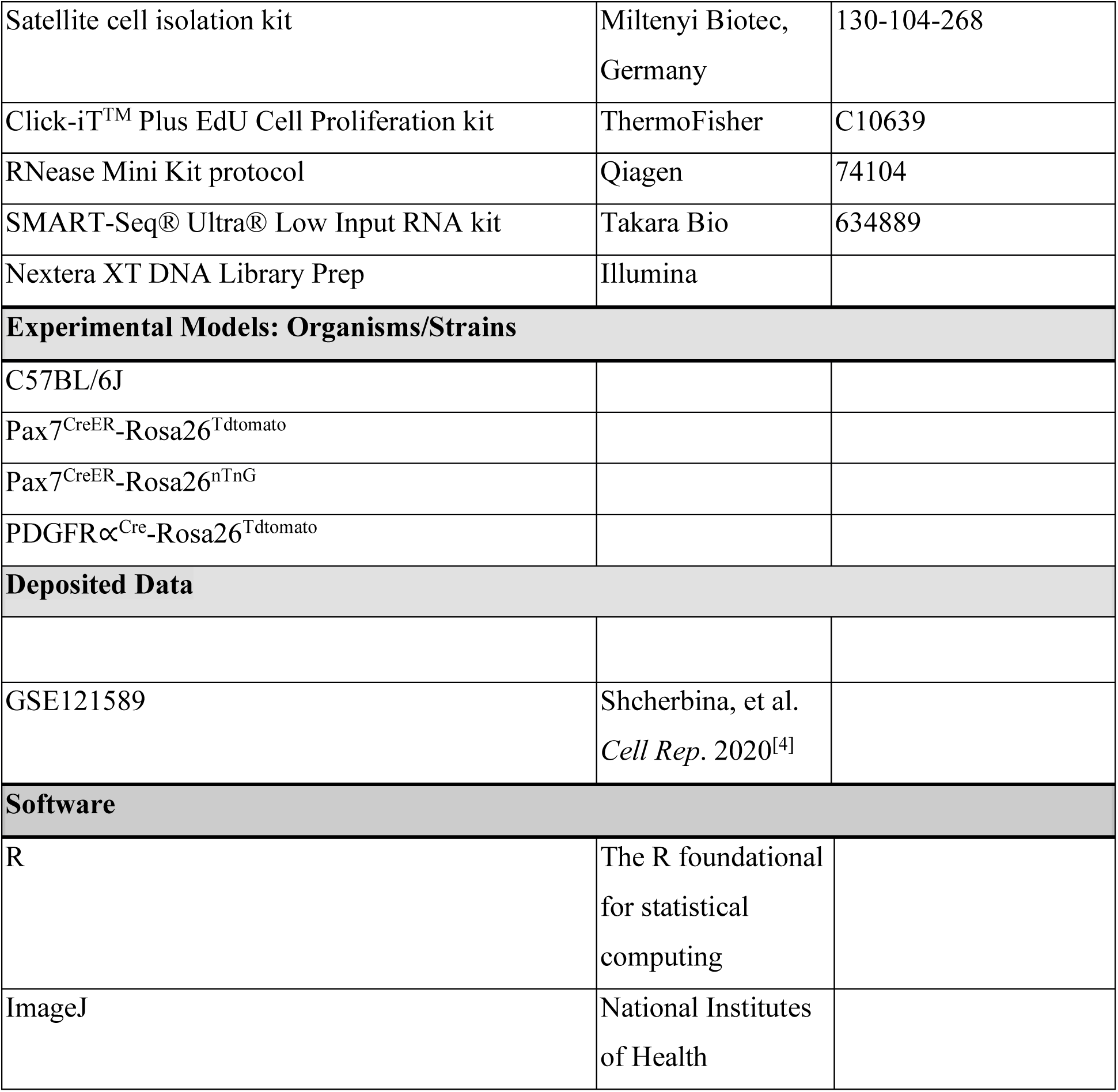

### Muscle Stem Cell (MuSC) Isolation via Magnetic-Activated Cell Sorting

Pax7^CreER^-Rosa26^Tdtomato^ (3 and 17-20 months old) and Pax7^CreER^-Rosa26^nTnG^ (13 and 24-26 months old) mice were injected with tamoxifen for 5 days and allowed to recover for 2 days. C57BL/6J wild-type mice were used for the co-staining of transcription factors (Pax7 and MyoD). Animals were sacrificed with CO_2_, and hindlimb muscles were harvested from both hindlimbs. Tissues were processed as previously described.^[62]^ Briefly, muscles were weighted, minced, transferred to 50 ml conical tubes with 20 ml of digest solution (2.5 U/ml Dispase II, 0.2% Collagenase II in Dulbecco’s Modified Eagle Medium per mouse) and incubated in a bead bath at 37°C for 1 hour. To break the muscle, pipetting of the tissue was performed every 30 minutes with an FBS coated pipette. Twenty ml of stop solution (20% heat inactivated FBS in F10 media) was then added to inactivate the enzymes. Digested muscle was filtered through a 70 µm cell strainer and centrifuged at 350xg for 5 min at 4°C. MuSCs were then isolated using the Satellite cell Isolation Kit following the manufacturer’s instructions. To further purify our MuSCs population, isolated cells were pre-plated in an uncoated tissue culture plate for 2 hours to remove excessive fibroblasts. ^[63]^ MuSCs were confirmed to be TdTomato^+^.

### Fibroadipogenic progenitors (FAPs) Isolation via Fluorescence-Activated Cell Sorting (FACs)

PDGFR∝^Cre^-Rosa26^Tdtomato^ mice (3 months old) were sacrificed with CO_2_, and hindlimb muscles were harvested and digested as described above. After filtration and centrifugation, cell suspensions were then rinsed with 4 ml of FACS buffer (2% of heat inactivated FBS in HBSS). The samples were then divided into tubes considering the controls (unstained, single colors and fluorescence minus one). Cells were then centrifuged at 350xg for 5 minutes at 4°C and supernatant was discarded. Samples were incubated with corresponding antibodies diluted in FACS media (CD45-2.5 µg/ml, CD31-5 µg/ml and SCa-1-0.5 µg/ml) for 30 minutes on a shaker plate. Then, 2-3 ml of FACS media was added to each tube and centrifuged (350xg minutes for 5 minutes at 4°C), supernatant was removed, and samples resuspended in FACS media. DAPI (1 µg/ml) was then added and samples were filtered through a 30 µm cell strainer and loaded into the Sony MA900 sorter. The gating strategy used to isolate Sca-1^+^ cells is shown on **Supp. Fig. 1C-F**. The negative (haematopoietic-CD45^+^ and endothelial-CD31^+^) and positive surface markers (stem cell antigen 1 or Sca-1^+^) were used to isolate FAPs,^[64]^ and FACS-isolated cells were confirmed to be TdTomato^+^.

### Culture of cells on Matrigel-coated coverslips

A 10% Matrigel solution in DMEM was added to coverslips (Fisherbrand, 12-545-80P) placed in a 24-non-treated well plate. After 2 minutes, the solution was removed, and the coverslips were incubated for 30 minutes at 37°C and allowed to dry for 1 hour at room temperature. C2C12s myoblasts, and isolated MuSCs and FAPs, were cultured on the Matrigel-coated coverslips at the corresponding cell density (MuSCs-7500 cells/cm^2^, FAPs-10000 cells/cm^2^, C2C12s-1000 cells/cm^2^) and incubated with their media (**Supp. Table 1**).

To understand the role of nascent matrix in the functionality of MuSCs, a reversible inhibitor of exocytosis was used to reduce nascent protein secretion. Exo-1 (120 nM) was added every day to affect the transport and secretion of proteins. At 5 days, fusion was induced by changing growth media (AHA-based) to differentiation media (5% horse serum and 1% pen/strep in DMEM). After 5 days, samples were fixed with 4% paraformaldehyde as described below.

### Preparation of Norbornene-modified hyaluronic acid (Nor-HA) hydrogels

Nor-HA polymer was synthesized as previously described.^[31]^ Nor-HA hydrogel precursor solution was prepared by dissolving Nor-HA polymer (2% wt/vol, 2 mg/100 µl) in PBS. Solution was then vortex, centrifuged and sonicated for 5 min. Photoinitiator (LAP) was then added at a 1:10 dilution, together with RGD peptide for cell adhesion (4µl/100µl, GCGYGRGDSPG), and cross-linker (DTT). The concentration of the former depended on the desired stiffness (Fig. S3A).

Methacrylated coverslips were prepared by immersing them for 3 minutes in methacrylated solution, which consisted of 1.5 ml of diluted acetic acid (1:10), 250 µl of methacrylate in 50 ml of Ethanol. Coverslips were then rinsed with Ethanol, and place on a dish with foil. Both sides of rectangle coverslips (50 mm, Fisherbrand, 12-545-GP) were immersed in sigmacote.

Six drops (20 µl each) of Nor-HA precursor solution were added to the sigmacote-treated slide, and the methacrylated coverslips were added on top with the methacrylated-treated side in contact with the solution. The sigmacote-treated slide with hydrogel solution and methacrylated coverslips was exposed to blue light (3 mW/cm^2^; EXFO OmniCure Series 2000) for 3 minutes. The hydrogels with coverslips were then detach carefully with forceps and place in a 24-non-treated well plates (Corning, 351147) with 300 µl of PBS. Hydrogels were then UV sterilized for 1 hour and PBS was replaced with sterile solution.

### 2D culture of MuSCs on Nor-HA hydrogels

Previously isolated MuSCs were counted with an hemacytometer and cultured on the hydrogels at a density of 7500 cells/ cm^2^. Methionine analog (AHA) based media was then added to the culture to label nascent proteins (supplementary table 1). To assess cellular proliferation, EdU was added to the cells at day 4 of culture and incubated for 24 hours.

For the growth factor supplementation, IGF-1 (25 ng/ml) was added to the AHA media, and the solution was changed every two days.

### Immunostaining and Imaging Analysis

To capture nascent matrix deposition, labeling of nascent proteins was done through a click chemistry reaction. AHA-based media was removed from the wells, and 300 µl of rinsing buffer (2% BSA in DPBS) was added for 2 minutes at room temperature (RT). Then, cells were incubated at 37°C for 35 minutes with 300 µl of AHA-labeling solution (45 µM of DBCO in rinsing buffer). Hydrogels were washed 3 x 2 minutes with the rinsing buffer, and cells were fixed with 300 µl of 4% paraformaldehyde for 30 minutes at RT. Samples were then rinsed 3 x 2 minutes with DPBS, followed by an incubation with 0.1% Cell Mask in DPBS for 30 minutes at RT to label the plasma membrane. Another 3 x 2 minutes rinses with DPBS were done, and samples were incubated with DAPI (1:1000) in DPBS for 20 minutes at RT, with 2 x5 minutes rinses in DPBS afterwards. Coverslips with hydrogels were then transferred to microscope slides and mounted with Pro-long Diamond anti-fade mountant.

For staining of vinculin and DDX6 with nascent proteins, samples were permeabilized (0.2% triton in DPBS) for 15 minutes after incubation with Cell Mask, rinsed 3x5 minutes with DPBS and blocked (2% BSA, 5% goat serum in DPBS) for 1 hour. Cells were then incubated either with vinculin (1:250) or DDX6 (1:1000) antibodies (diluted in 1%BSA in DBST) overnight at 4°C. Cells were rinsed 3 x 3 minutes in DPBST (0.1% tween-20) and incubated for 2 hours at RT with secondary antibody (Alexa Fluor-647 goat anti-mouse IgG-1:500 (vinculin), 1:1000 (DDX6), 1%BSA in DBST). Samples were then rinsed with DPBST, incubated with DAPI, washed again and mounted as described above. The co-staining of Pax7 and MyoD with nascent proteins was done as described above. Cells were incubated with blocking buffer (2% BSA, 5% goat serum, 0.1% tween in DPBS) for 1 hour, followed by primary antibody incubation (Pax7-1:100, MyoD-1:100, 1%BSA in DBST) overnight at 4°C. Cells were rinsed 3 x 3 minutes in DPBST and incubated for 2 hours at RT with secondary antibodies (Alexa Fluor-555 goat anti-mouse IgG1-1:200 and Alexa Fluor-594 goat anti-mouse IgG2b-1:200, 1%BSA in DBST). EdU labeling was done following the instructions of the manufacturer. Lastly, fixed myotubes were blocked for 1 hour (1% BSA, 22.52 mg/ml glycine in DPBST) and incubated with Myh3 antibody (1:20, 1%BSA in DBST) overnight at 4°C. After rinsing with DPBST, cells were incubated for 2 hours at RT with secondary antibody (Alexa Fluor-555 goat anti-mouse IgG1-1:100).

For the analysis of nascent matrix alone, Z-stacks were obtained with the Nikon A1si Confocal Inverted, using a 60X oil-immersion objective. For co-staining with transcription factors, EdU, and DDX6, imaging was done with the Leica Stellaris 5 Confocal Inverted, using a 63X oil-immersion objective. All images were scanned with the same imaging parameters for each microscope. Z-projection using maximum intensity was obtained for each channel using ImageJ software. Maximum projection images were then used to quantify total area of single MuSC nascent matrix deposition. Binary image of CellMask channel was subtracted from the DBCO channel to avoid quantification of intracellular proteins. Analyze particle function was then utilized to quantify total area for nascent matrix. Pax7 and MyoD intensities were quantified with the mean fluorescence intensity measurement in ImageJ. Binary images of DDX6 were acquired and analyze particle function was used to quantify the foci per cell. Imaging of myotubes was done with a 10X objective using the Nikon microscope. The fusion index was quantified as the total number of nuclei per field of view divided by the number of nuclei in myotubes. For the myotube width, 5-10 measurements were done along the myotube.

### mRNA sequencing

To determine mechanisms related to the impaired nascent matrix, MuSCs from young and old mice were isolated via magnetic activated cell sorting and seeded on the norbornene-hyaluronic acid hydrogels at a cell density of 7500 cells/ cm^2^. After 5 days on culture, cells were trypsinized and RNA was isolated following the RNease Mini Kit protocol. cDNAs were prepared based on the SMART-Seq® Ultra® Low Input RNA Kit, and libraries prepared following the Nextera XT DNA Library Prep protocol. Libraries were sequenced on NovaSeq instrument (Illumina) with paired-end reads and 25 million reads per sample.

### RNA-Seq Data Processing and Analysis

The sequencing reads were demultiplexed into FASTQ files and the adapters were trimmed using the Cutadapt v2.3. The reads were mapped to the mouse genome using Kallisto.^[65]^ Counts were then assigned to genes using RSEM. ^[66]^Differential expression analysis of old versus young groups was performed with DESeq2^[67]^ and significantly differentially expressed genes were obtained with a p adjusted value<0.05. We then used WebGestalt^[68]^ to perform a Gene Set Enrichment Analysis with the Reactome pathway database.^[40]^ Differentially expressed pathways were visualized with a bubble plot indicating the false discovery rate and the gene number.

For previously obtained RNA-sequencing data^[4]^, sequencing reads were analyzed as mentioned above. Differential expression analysis between each time point after injection versus uninjured group was performed with DESeq2. Significantly differentially expressed genes (p adjusted value<0.05) were segregated to include only ECM-related genes (core matrisome) based on a previous classification^[31, 35]^ and visualized with a heatmap.

### Statistical Analysis

Data (mean±SEM) were analyzed by student’s t-test, and one-or two-way ANOVA with Tukey’s post hoc pairwise comparisons with significance set at a p value of 0.05. Graph Pad Prism 9 (version 9.1.1) was used for the statistical analysis.

## Acknowledgments

Research reported in this publication was partially supported by the University of Michigan Geriatrics Center and National Institute of Aging under award number P30 AG024824 (PD, CAA), Genentech Research Award (CAA), the 3M Foundation (CAA), the Department of Defense and Congressionally Directed Medical Research Program W81XWH2010336 and W81XWH2110491 (CAA), a National Science Foundation CAREER award (2045977), Defense Advanced Research Projects Agency (DARPA) “BETR” award D20AC0002 (CAA) awarded by the U.S. Department of the Interior (DOI), Interior Business Center, and the Hevolution HF-AGE award (CAA). The content is solely the responsibility of the authors and does not necessarily represent the official views of the National Institutes of Health or National Science Foundation, the position or the policy of the Government, and no official endorsement should be inferred.

## Data availability Statement

Data associated with this study are available in the main text or the supplementary materials. Bulk RNA-sequencing data are available from the NCBI GEO database with the accession number GSE252829.

## Author contributions

P.D., E.P., and M.E. performed experiments. B.A.Y.. analyzed data. P.D., C.L. and C.A.A. designed the experiments. P.D. and C.A.A. wrote the manuscript with additions from other authors.

---

**Scheme 1.**
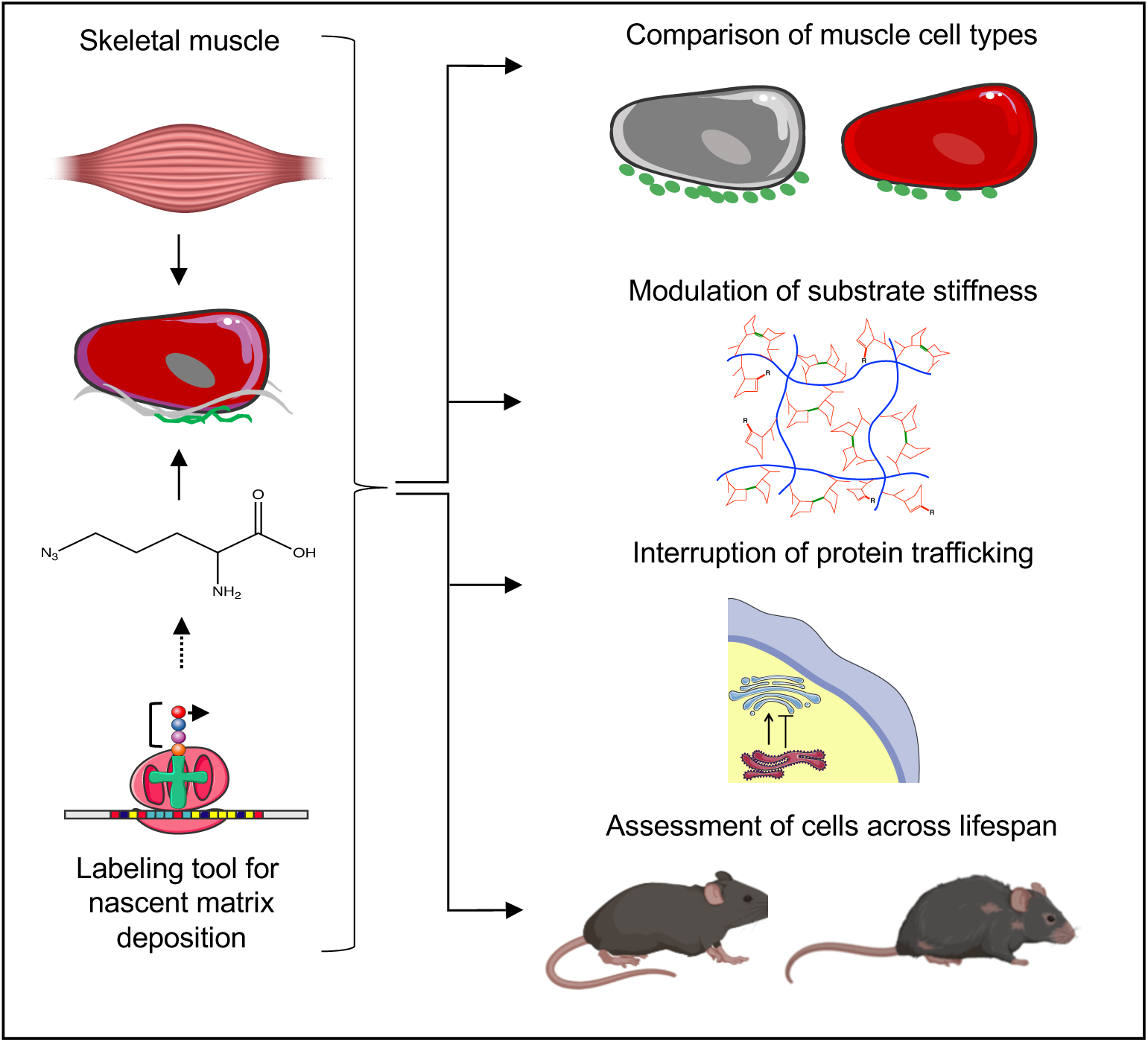
Quantification and comparison of nascent matrix deposition from muscle stem cells (MuSCs). MuSCs were contrasted in different conditions to explore how these cells deposit nascent matrix and remodel their niche. Comparisons were made with other cell types (fibro-adipogenic progenitors), on different physiological muscle stiffnesses matching homeostasis and injury, blocking protein trafficking and across lifespan. These results show direct measurements of how adult stem cells remodel their niche.

## References

1. E. Fuchs and H.M. Blau, Cell Stem Cell. 2020, 27, 532.

2. N.A. Dumont, Y.X. Wang, and M.A. Rudnicki, Development. 2015, 142, 1572.

3. M. Loreti and A. Sacco, NPJ Regen Med. 2022, 7, 16.

4. A. Shcherbina, J. Larouche, P. Fraczek, B.A. Yang, L.A. Brown, J.F. Markworth, C.H. Chung, M. Khaliq, K. de Silva, J.J. Choi, M. Fallahi-Sichani, S. Chandrasekaran, Y.C. Jang, S.V. Brooks, and C.A. Aguilar, Cell Rep. 2020, 32, 107964.

5. L. Liu, T.H. Cheung, G.W. Charville, B.M. Hurgo, T. Leavitt, J. Shih, A. Brunet, and T.A. Rando, Cell Rep. 2013, 4, 189.

6. B.A. Yang, J.A. Larouche, K.M. Sabin, P.M. Fraczek, S.C.J. Parker, and C.A. Aguilar, Aging Cell. 2023, 22, e13789.

7. J.V. Kurland, A.A. Cutler, J.T. Stanley, N.D. Betta, A. Van Deusen, B. Pawlikowski, M. Hall, T. Antwine, A. Russell, M.A. Allen, R. Dowell, and B. Olwin, Stem Cell Reports. 2023, 18, 1325.

8. J.C. Kimmel, N. Yi, M. Roy, D.G. Hendrickson, and D.R. Kelley, Cell Rep. 2021, 35, 109046.

9. L. Lukjanenko, S. Karaz, P. Stuelsatz, U. Gurriaran-Rodriguez, J. Michaud, G. Dammone, F. Sizzano, O. Mashinchian, S. Ancel, E. Migliavacca, S. Liot, G. Jacot, S. Metairon, F. Raymond, P. Descombes, A. Palini, B. Chazaud, M.A. Rudnicki, C.F. Bentzinger, and J.N. Feige, Cell Stem Cell. 2019, 24, 433.

10. A.B. Hwang and A.S. Brack, Curr Top Dev Biol. 2018, 126, 299.

11. S.C. Schuler, J.M. Kirkpatrick, M. Schmidt, D. Santinha, P. Koch, S. Di Sanzo, E. Cirri, M. Hemberg, A. Ori, and J. von Maltzahn, Cell Rep. 2021, 35, 109223.

12. J.V. Chakkalakal, K.M. Jones, M.A. Basson, and A.S. Brack, Nature. 2012, 490, 355.

13. L. Lukjanenko, M.J. Jung, N. Hegde, C. Perruisseau-Carrier, E. Migliavacca, M. Rozo, S. Karaz, G. Jacot, M. Schmidt, L. Li, S. Metairon, F. Raymond, U. Lee, F. Sizzano, D.H. Wilson, N.A. Dumont, A. Palini, R. Fassler, P. Steiner, P. Descombes, M.A. Rudnicki, C.M. Fan, J. von Maltzahn, J.N. Feige, and C.F. Bentzinger, Nat Med. 2016, 22, 897.

14. L.K. Wood, E. Kayupov, J.P. Gumucio, C.L. Mendias, D.R. Claflin, and S.V. Brooks, J Appl Physiol (1985). 2014, 117, 363.

15. C. Rosant, M.D. Nagel, and C. Perot, Exp Gerontol. 2007, 42, 301.

16. J.M. Haus, J.A. Carrithers, S.W. Trappe, and T.A. Trappe, J Appl Physiol (1985). 2007, 103, 2068.

17. J. Etienne, C. Liu, C.M. Skinner, M.J. Conboy, and I.M. Conboy, Skelet Muscle. 2020, 10, 4.

18. M. Rozo, L. Li, and C.M. Fan, Nat Med. 2016, 22, 889.

19. A.S. Brack, M.J. Conboy, S. Roy, M. Lee, C.J. Kuo, C. Keller, and T.A. Rando, Science. 2007, 317, 807.

20. I.M. Conboy, M.J. Conboy, A.J. Wagers, E.R. Girma, I.L. Weissman, and T.A. Rando, Nature. 2005, 433, 760.

21. S.S. Rayagiri, D. Ranaldi, A. Raven, N.I.F. Mohamad Azhar, O. Lefebvre, P.S. Zammit, and A.G. Borycki, Nat Commun. 2018, 9, 1075.

22. J.S. Silver, K.A. Gunay, A.A. Cutler, T.O. Vogler, T.E. Brown, B.T. Pawlikowski, O.J. Bednarski, K.L. Bannister, C.J. Rogowski, A.G. McKay, F.W. DelRio, B.B. Olwin, and K.S. Anseth, Sci Adv. 2021, 7,

23. P.M. Gilbert, K.L. Havenstrite, K.E. Magnusson, A. Sacco, N.A. Leonardi, P. Kraft, N.K. Nguyen, S. Thrun, M.P. Lutolf, and H.M. Blau, Science. 2010, 329, 1078.

24. C.M. Madl, I.A. Flaig, C.A. Holbrook, Y.X. Wang, and H.M. Blau, Biomaterials. 2021, 275, 120973.

25. P.M. Gilbert, S. Corbel, R. Doyonnas, K. Havenstrite, K.E. Magnusson, and H.M. Blau, Integr Biol (Camb). 2012, 4, 360.

26. W.M. Han, S.E. Anderson, M. Mohiuddin, D. Barros, S.A. Nakhai, E. Shin, I.F. Amaral, A.P. Pego, A.J. Garcia, and Y.C. Jang, Sci Adv. 2018, 4, eaar4008.

27. M.T. Tierney, A. Gromova, F.B. Sesillo, D. Sala, C. Spenle, G. Orend, and A. Sacco, Cell Rep. 2016, 14, 1940.

28. M.M. Stern, R.L. Myers, N. Hammam, K.A. Stern, D. Eberli, S.B. Kritchevsky, S. Soker, and M. Van Dyke, Biomaterials. 2009, 30, 2393.

29. H. Yi, S. Forsythe, Y. He, Q. Liu, G. Xiong, S. Wei, G. Li, A. Atala, A. Skardal, and Y. Zhang, Acta Biomater. 2017, 62, 222.

30. K.J. Wilschut, H.P. Haagsman, and B.A. Roelen, Exp Cell Res. 2010, 316, 341.

31. C. Loebel, A.M. Saleh, K.R. Jacobson, R. Daniels, R.L. Mauck, S. Calve, and J.A. Burdick, Nat Protoc. 2022, 17, 618.

32. K. Thomas, A.J. Engler, and G.A. Meyer, Connect Tissue Res. 2015, 56, 1.

33. B. Biferali, D. Proietti, C. Mozzetta, and L. Madaro, Front Physiol. 2019, 10, 1074.

34. C.A. Aguilar, R. Pop, A. Shcherbina, A. Watts, R.W. Matheny, Jr., D. Cacchiarelli, W.M. Han, E. Shin, S.A. Nakhai, Y.C. Jang, C.T. Carrigan, C.A. Gifford, M.A. Kottke, M. Cesana, J. Lee, M.L. Urso, and A. Meissner, Stem Cell Reports. 2016, 7, 983.

35. A.M. Saleh, K.R. Jacobson, T.L. Kinzer-Ursem, and S. Calve, Cell Mol Bioeng. 2019, 12, 495.

36. Y. Feng, S. Yu, T.K. Lasell, A.P. Jadhav, E. Macia, P. Chardin, P. Melancon, M. Roth, T. Mitchison, and T. Kirchhausen, Proc Natl Acad Sci U S A. 2003, 100, 6469.

37. K. Mishev, W. Dejonghe, and E. Russinova, Chem Biol. 2013, 20, 475.

38. P. Sousa-Victor, L. Garcia-Prat, and P. Munoz-Canoves, Nat Rev Mol Cell Biol. 2022, 23, 204.

39. S.C. Schuler, N. Gebert, and A. Ori, Mech Ageing Dev. 2020, 190, 111288.

40. A. Fabregat, K. Sidiropoulos, G. Viteri, O. Forner, P. Marin-Garcia, V. Arnau, P. D’Eustachio, L. Stein, and H. Hermjakob, BMC Bioinformatics. 2017, 18, 142.

41. R.C. Taylor and A. Dillin, Cold Spring Harb Perspect Biol. 2011, 3,

42. C. Guillet and Y. Boirie, Diabetes Metab. 2005, 31 Spec No 2, 5S20.

43. A. Philippou and E.R. Barton, Growth Horm IGF Res. 2014, 24, 157.

44. B.A. Hemmings and D.F. Restuccia, Cold Spring Harb Perspect Biol. 2015, 7,

45. S. Park, D.G. Gonzalez, B. Guirao, J.D. Boucher, K. Cockburn, E.D. Marsh, K.R. Mesa, S. Brown, P. Rompolas, A.M. Haberman, Y. Bellaiche, and V. Greco, Nat Cell Biol. 2017, 19, 155.

46. M. Adler, N. Moriel, A. Goeva, I. Avraham-Davidi, S. Mages, T.S. Adams, N. Kaminski, E.Z. Macosko, A. Regev, R. Medzhitov, and M. Nitzan, Cell Rep. 2023, 42, 112412.

47. M.N. Wosczyna, C.T. Konishi, E.E. Perez Carbajal, T.T. Wang, R.A. Walsh, Q. Gan, M.W. Wagner, and T.A. Rando, Cell Rep. 2019, 27, 2029.

48. J.A. Larouche, P.M. Fraczek, S.J. Kurpiers, B.A. Yang, C. Davis, J.A. Castor-Macias, K. Sabin, S. Anderson, J. Harrer, M. Hall, S.V. Brooks, Y.C. Jang, N. Willett, L.D. Shea, and C.A. Aguilar, Proc Natl Acad Sci U S A. 2022, 119, e2111445119.

49. J.A. Larouche, E.C. Wallace, B.D. Spence, E. Buras, and C.A. Aguilar, JCI Insight. 2023, 8,

50. P. Duran, F. Boscolo Sesillo, M. Cook, L. Burnett, S.A. Menefee, E. Do, S. French, G. Zazueta-Damian, M. Dzieciatkowska, A.J. Saviola, M.M. Shah, C. Sanvictores, K.G. Osborn, K.C. Hansen, M. Shtrahman, K.L. Christman, and M. Alperin, Sci Transl Med. 2023, 15, eabj3138.

51. J. Wang, A. Khodabukus, L. Rao, K. Vandusen, N. Abutaleb, and N. Bursac, Biomaterials. 2019, 221, 119416.

52. B.A. Yang, T.M. Westerhof, K. Sabin, S.D. Merajver, and C.A. Aguilar, Adv Sci (Weinh). 2021, 8, 2002825.

53. M.T. Webster, U. Manor, J. Lippincott-Schwartz, and C.M. Fan, Cell Stem Cell. 2016, 18, 243.

54. E. Jacques, Y. Kuang, A.P. Kann, F. Le Grand, R.S. Krauss, and P.M. Gilbert, Elife. 2022, 11,

55. C.M. Madl, B.L. LeSavage, R.E. Dewi, K.J. Lampe, and S.C. Heilshorn, Adv Sci (Weinh). 2019, 6, 1801716.

56. P. Shi, K. Shen, S. Ghassemi, J. Hone, and L.C. Kam, Cell Mol Bioeng. 2009, 2, 464.

57. G. Dura, M. Crespo-Cuadrado, H. Waller, D.T. Peters, A. Ferreira-Duarte, J.H. Lakey, and D.A. Fulton, Macromol Biosci. 2022, 22, e2200134.

58. M. Quarta, J.O. Brett, R. DiMarco, A. De Morree, S.C. Boutet, R. Chacon, M.C. Gibbons, V.A. Garcia, J. Su, J.B. Shrager, S. Heilshorn, and T.A. Rando, Nat Biotechnol. 2016, 34, 752.

59. S.Y. Yang, M. Hoy, B. Fuller, K.M. Sales, A.M. Seifalian, and M.C. Winslet, Lab Invest. 2010, 90, 391.

60. T. Yoshida and P. Delafontaine, Cells. 2020, 9,

61. Y. Kanazawa, R. Miyachi, T. Higuchi, and H. Sato, Int J Mol Sci. 2023, 24,

62. J.A. Larouche, M. Mohiuddin, J.J. Choi, P.J. Ulintz, P. Fraczek, K. Sabin, S. Pitchiaya, S.J. Kurpiers, J. Castor-Macias, W. Liu, R.L. Hastings, L.A. Brown, J.F. Markworth, K. De Silva, B. Levi, S.D. Merajver, G. Valdez, J.V. Chakkalakal, Y.C. Jang, S.V. Brooks, and C.A. Aguilar, Elife. 2021, 10,

63. K. Yoshioka, Y. Kitajima, N. Okazaki, K. Chiba, A. Yonekura, and Y. Ono, Front Cell Dev Biol. 2020, 8, 793.

64. A.W. Joe, L. Yi, A. Natarajan, F. Le Grand, L. So, J. Wang, M.A. Rudnicki, and F.M. Rossi, Nat Cell Biol. 2010, 12, 153.

65. N.L. Bray, H. Pimentel, P. Melsted, and L. Pachter, Nat Biotechnol. 2016, 34, 525.

66. B. Li and C.N. Dewey, BMC Bioinformatics. 2011, 12, 323.

67. M.I. Love, W. Huber, and S. Anders, Genome Biol. 2014, 15, 550.

68. B. Zhang, S. Kirov, and J. Snoddy, Nucleic Acids Res. 2005, 33, W741.

