## Supplementary Material for "Quantification of local matrix deposition during muscle stem cell activation using engineered hydrogels"

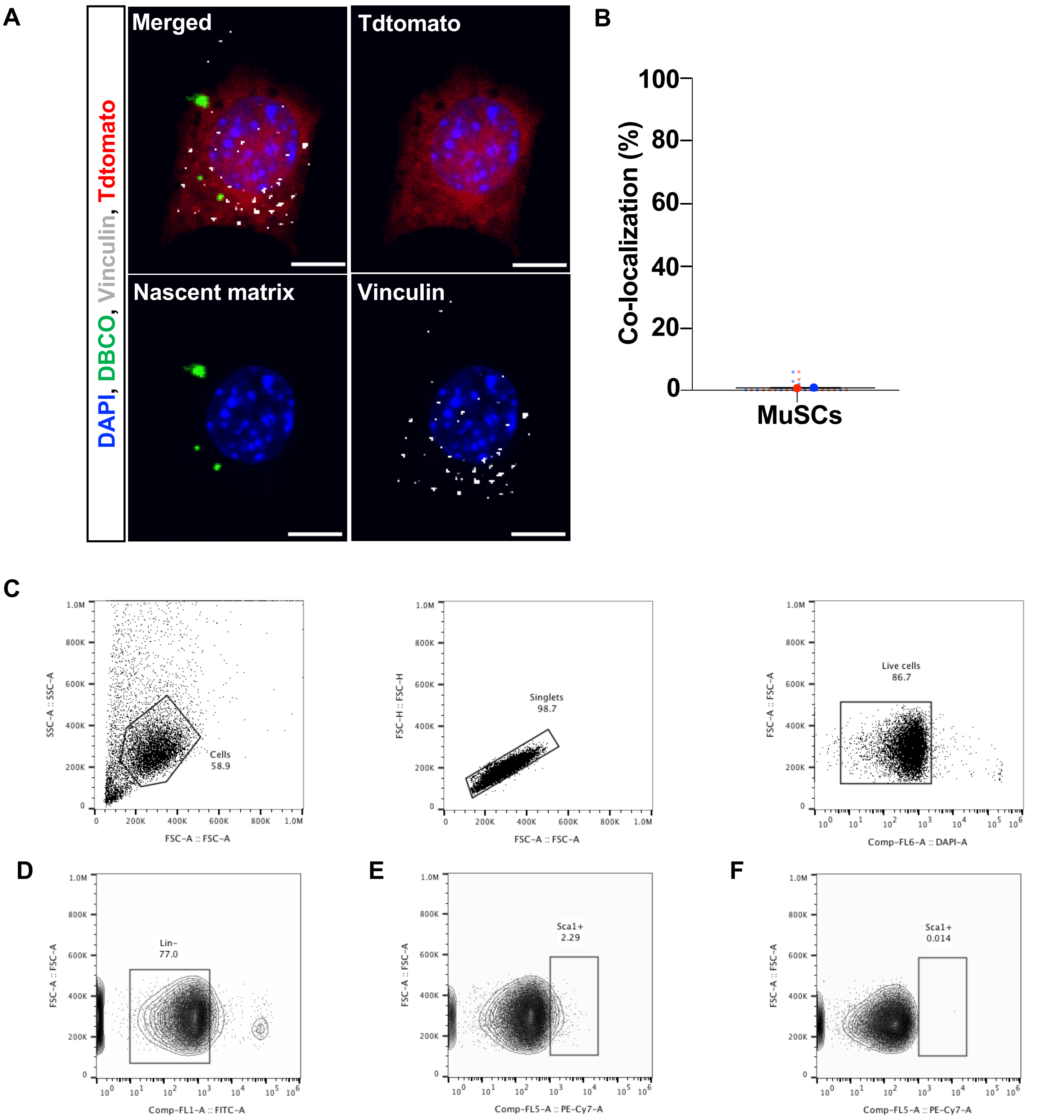


**Supplemental Figure 1. A)** Representative image of muscle stem cell stained for nascent proteins and vinculin, a focal adhesion protein. Blue: DAPI, Green: Nascent Matrix, Gray: Vinculin, Red: TdTomato. Scale = 5 $\mu$m. **B)** Quantification of the colocalization of the area of nascent matrix with vinculin. Small circles represent single cells (n=12/replicate), and large circles indicate the biological replicates (n=2 biological replicates). Representative FACS plots showing gating for live cells **(C)**, lineage negative (CD-31, CD45, and Ter-119, **D**) and positive (Sca-1, **E**) surface markers to isolate fibro-adipogenic progenitors. Numbers within gates indicate percentage of cells within gate. **F)** Example of negative control.

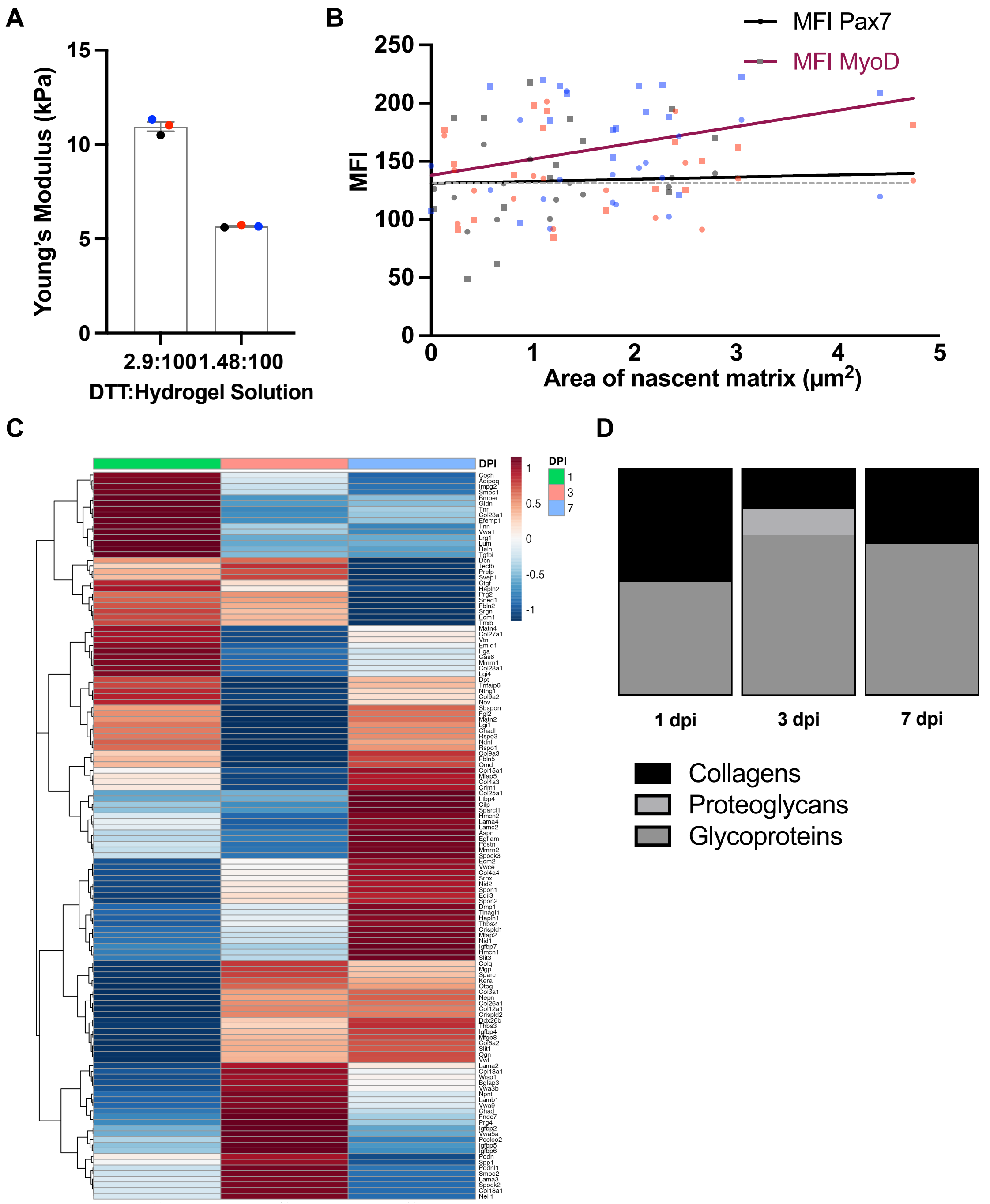


**Supplemental Figure 2. A)** Young's modulus of norbornene-modified hyaluronic acid (Nor-HA) hydrogel prepared with different concentration of cross-linker (DTT). Biological replicates=3. **B)** Linear regression of the area of nascent matrix with the mean fluorescence intensity (MFI) of Pax7 and MyoD of MuSCs cultured on 11 kPa Nor-HA hydrogel. Single cells=10-25/replicates. **C)** Heatmap of differentially expressed ECM-related genes (core matrisome) in MuSCs at 1, 3, and 7 days after barium chloride injection (GSE121589). **D)** Proportion of the significantly upregulated genes from the core matrisome categories at each timepoint after injury.

**A**

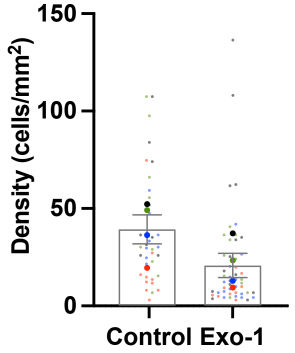


**Supplemental Figure 3. A)** Quantification of cell density with and without Exo-1. mean ± SEM; small circles represent single cells (n=10-30/replicate) and large circles indicate the biological replicates (N=4).

A

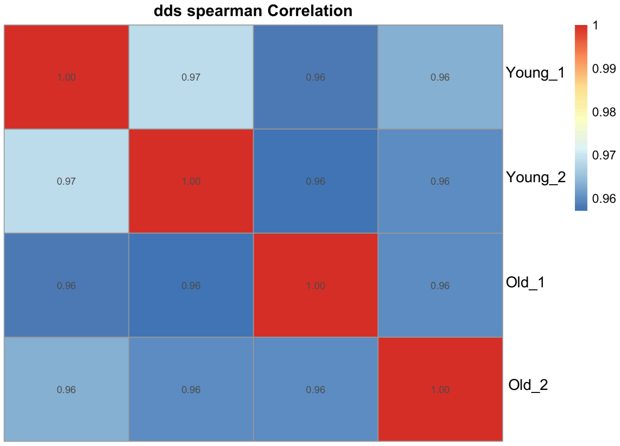


**Supplemental Figure 4. A)** Spearman correlation coefficient of the gene expression profiles for the biological replicates isolated from young and old muscle stem cells.

**Supplementary Table 1. Cell culture media used to label nascent proteins.**

| | | **Muscle stem cells** | **Fibroadipogenic progenitors** | **C2C12s** |
|---|---|---|---|---|
| AHA-based stock media | DMEM (free sodium pyruvate, glutamax, and methionine) | 86.7% | 81.8% | 81.8% |
| | L-methionine | 0% | 0.15% | 0.015% |
| | L-cystine | 0.096% | 0.096% | 0.096% |
| | Glutamax | 1% | 1% | 1% |
| | Sodium pyruvate | 0.9% | 0.9% | 0.9% |
| | AHA | 0.2% | 0.1% | 0.1% |
| | Ascorbic acid | 0.1% | 0.1% | 0.1% |
| | FBS | 10% | 15% | 15% |
| | Pen-strep | 1% | 1% | 1% |
| | AHA-based stock media | 50% | | |
| | F-10 | 40% | | |
| | FBS | 9.5% | | |
| | Pen-strep | 0.2% | | |
| | bFGF | 0.02 µg/ml | 2.5 ng/ml | |
